# Coevolution of Codependent Hosts and Symbionts

**DOI:** 10.64898/2026.07.21.739856

**Authors:** Michael Lynch, Kunaal Joshi, Adrian González-Casanova

**Affiliations:** Biodesign Center for Mechanisms of Evolution, Arizona State University, Tempe, AZ 85287

## Abstract

Many endosymbioses in eukaryotes superficially appear to be beneficial to both participants. However, there is little direct evidence for this, and symbioses naturally set up conditions in which each member of the pair is under selection to extract resources from the other. Ultimately, the endosymbiont either evolves to be in conflict with the interests of the host or to act cooperatively with the host contrary to its own best interests. Focusing on obligate symbioses, we develop theory to clarify the population-genetic conditions favoring the alternative outcomes. The balance is usually tipped in favor of exploitation by the symbiont, particularly when the number of symbionts within host cells is high, selection is strong on symbionts relative to hosts, there is horizontal transfer of symbionts, and/or the symbionts have accelerated mutation rates or turnover times. If the symbiont conditions the host-cell biology to enhance within-host population sizes, selection for selfish symbionts will be further enhanced by the diminished level of within-host drift. Although the host evolves in parallel to exploit resources from the endosymbiont, the net result is often a stalemate in which the host is no better off than prior to host-symbiont coevolution. Strict vertical inheritance can result in an evolutionary alignment of interests of the endosymbiont and the host, as this minimizes the possibility of within-host selection, but even here there is a critical host population size below which the symbiont evolves to exploit the host. These results suggest that the evolutionary enslavement of a symbiont to benefit a host species requires a narrow mix of population-biological features of both participants.

---

Although the influence of symbioses on community structure has been appreciated for well over a century, relatively little is known about the evolutionary paths down which such interactions proceed once established. Extreme forms of symbiosis range from parasitism, whereby one member of the pair exploits the other, to mutualisms in which both members are thought to obtain advantages. However, even the latter condition can often be an illusion, in the sense that while both participants have become codependent, one or both might still perform no better or even worse than had been the case prior to establishment of the interaction. The latter condition is sometimes called addictive symbiosis (Castelli et al. 2024; Hammer 2024), a key question being whether what superficially appears to be mutually beneficial actually involves exploitation (Bronstein 1994).

Symbioses can be facultative or obligatory, with the latter referring to situations in which one or both participants are incapable of independent living. Although the former is much more common and certainly of interest, deciphering general evolutionary consequences of facultative interactions is complicated by the need to include matters involving association and re-assortment, fitnesses of individuals with and without symbionts, the involvement of third-party species, etc. Thus, given the complexities that can emerge with even the simplest of systems, the focus here is on obligate symbioses, with a goal of developing general models for alternative evolutionary outcomes in different selection and population-genetic settings. Well-known obligate symbioses include the amino-acid providing endosymbiotic bacteria found in numerous sap-feeding insects (Moran 1996; Moran and Plague 2004; McCutcheon and Von Dohlen 2011; McCutcheon et al. 2019), the photosynthetic organelle of the amoeboid protist *Paulinella* (Gabr et al. 2020), the nitroplasts found in various marine phytoplankters (Coale et al. 2024; Verma and White 2025), and the killer bacteria of *Paramecium* (Görtz and Fokin 2009). In addition, many bacteriophage and plasmids ensure their long-term maintenance within bacterial host cells by deploying toxin-antitoxin systems (Gao et al. 2020; Mendoza-Guido and Rojas-Jimenez 2025).

Obligate symbioses are also relevant to the establishment of cell-biological features at deeper points in the history of life. For example, it has been suggested that the proto-ribosome started out as a molecular parasite or mutualist (Krupovic and Koonin 2026; Lynch and Ellington 2026). Given its phylogenetic affinity with intracellular pathogenic bacteria and the reduced bioenergetic capacity of host eukaryotic cells, the mitochondrion might have also started as an intracellular parasite (Lynch 2024). The spliceosome, the eukaryotic machine used to remove introns from precursor mRNAs, almost certainly originated as a parasitic Group-II intron (Lynch 2007). In each of these cases, what may have started out as a deleterious interaction from the standpoint of the host cell would have become permanently stabilized once each member of the pair relinquished a key function to the other. For example, once membrane bioenergetics were relocated from the cell membrane to the mitochondrion and genes from the mitochondrion were relocated to the host-cell nucleus, host and endosymbiont were reciprocally preserved. Likewise, once the the ancestral eukaryotic genome became populated with introns, the spliceosome was permanently endowed with a mandatory cell function. The establishment of such codependencies is conceptually similar to the subfunctionalization process that underlies the preservation of duplicate genes by degenerative mutations (Force et al. 1999; Lynch et al. 2001). All of these observations raise questions about the degree to which obligate mutualisms, often perceived as being reciprocally beneficial, actually represent energetic advances by one or both participants, i.e., which members of the pair gain advantages at the expense of the other.

The evolution of symbioses involves two levels of selection. At the endosymbiont level, selection favors mutants that proliferate at the highest rates within hosts cells. In contrast, host genotypes are selected on the basis of the joint effects of their own encoded traits and those of the symbionts that they harbor. Here, we develop a quantitative-genetic model for the general fitness features of obligatory two-species symbioses, exploring how the long-term evolutionary outcome depends on the basic population-genetic parameters of the participants (mutation rates and population sizes), the level of exchange of symbionts among hosts by horizontal transfer, and the magnitude of the strength of selection operating on the two species. As will be seen below, subtle changes in the population-genetic features of the two species may lead the system to evolve in the direction of either cooperation or conflict, i.e., the endosymbiont may evolve to either provision or extract resources from the host.

Many theoretical studies have considered the evolution of symbioses. However, most such studies have been concerned with the establishment and long-term stability of facultative vs. obligatory involvement of one or both partners (Nguyen and van Baalen 2001; Foster and Wenseleers 2006; Athreya et al. 2025; Prigent and Mullon 2026). For mathematical convenience, these studies often assume low or zero levels of mutation, a condition that will generally not be met in microbial systems, and deterministic population dynamics, a condition that never exists for newly arisen mutations (Doebeli and Knowlton 1998). Often, the genetic features of evolution do not enter the theory, with the primary focus being on ecological time scales starting with fixed sets of interactions (e.g., Sieber et al. 2021; Cosmo et al. 2023; Piñero et al. 2025; Ruiz-Herrera 2025). Various models have been developed for the evolution of organelle variants, including those involving two levels of selection (Bergstrom and Pritchard 1998; Roze et al. 2005; Edwards et al. 2021), but in these studies the host is generally assumed to be evolutionarily static, and/or the focus is on deleterious-mutation accumulation.

Although it is commonly thought that a stable symbiosis requires the alignment of interests between the two members of the pair (Frank 1983; Maynard Smith and Szathmary 1997; Michod 2000; West et al. 2015), the following theory suggests that this is not generally the case, demonstrating that the population-genetic details can greatly influence the evolutionary fates of both members of the pair. Here, we evaluate the long-term steady-state features of composite genotypes of host and symbiont partners experiencing arbitrary levels of selection, mutation, migration, and random genetic drift. We start with a relatively simple model in which hosts and symbionts have a range of stable population sizes and equal generation times, and then examine the situation in which the within-host symbiont population size is influenced by host-cell conditioning. Finally, we address the evolutionary consequences of different generation lengths of the participating partners.

## Characterization of the Model

We start with the assumption that the genotypes of host cells and their obligate endosymbionts (hereafter abbreviated to symbionts) can be characterized by composite quantitative traits, treated as integers centered around 0, and denoted as *h*_*i*_ with lower and upper bounds of −*E*_*H*_ and +*E*_*H*_ for the host (*H*), and as *s*_*j*_ with bounds of −*E*_*S*_ and +*E*_*S*_ for the symbiont (*S*) (Figure 1). These composite measures can be viewed in a variety of ways, ranging from the summed effects of mutations at individual nucleotide sites within single genetic loci in the two partners to the summed effects over alternative genotypes at different loci. The main assumption is that effects are additive across sites / loci on an underlying phenotypic scale.

**Figure 1.**
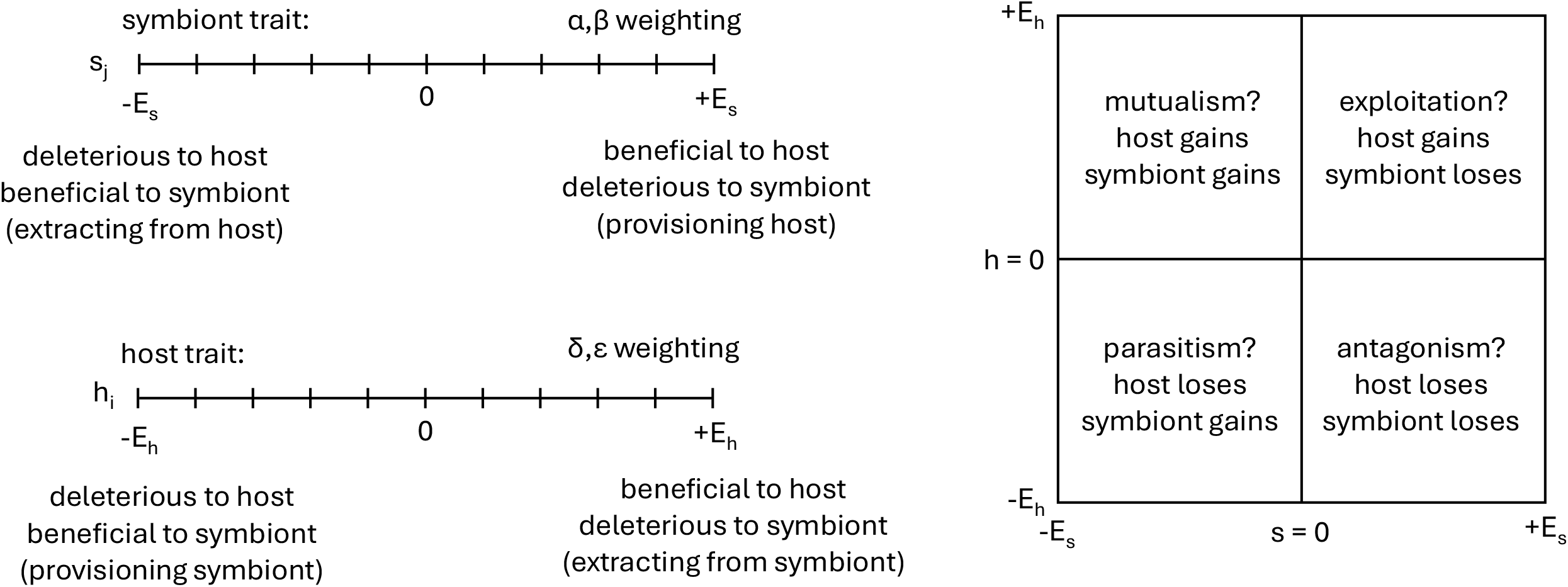
**Left)** Schematic for the underlying phenotypic traits of host and symbiont and their associated fitness functions. **Right)** Interpretations of potential evolutionary outcomes; the precise outcomes depend on the relative strengths of three selection coefficients.

The total genetic values on these scales are obtained by treating the *L*_*x*_ = 2*E*_*x*_ sites as biallelic (+/-), so that with *i* and *j* denoting the numbers of + alleles in the host and symbiont genomes,

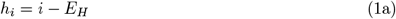

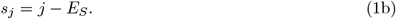

There are thus (*L*_*x*_ + 1) possible haplotypes for both host and symbiont genomes, which we assume to be haploid and asexual, although reassortment of symbionts via horizontal transfer among host cells is allowed (as discussed below).

The fitness of host-cell type *i* containing symbiont type *j* is denoted as

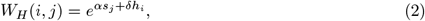

which yields *W*_*H*_ (0, 0) = 1 when *s*_*j*_ = *h*_*i*_ = 0 (at the midpoint of the phenotypic arrays) and when *αs*_*j*_ = −*δh*_*i*_. Under this exponential fitness function, *α* and *δ* denote selective advantages of incremental increases of *s*_*j*_ and *h*_*i*_, respectively, with positive values of the host and symbiont traits having beneficial effects on host fitness, and negative values diminishing host fitness. In other words, positive *s*_*j*_ implies a cooperative symbiont that provisions the host, and positive *h*_*i*_ implies a host’s additional capacity for extracting resources from the symbiont.

In contrast, the fitness of symbiont type *j* within host-cell type *i* is denoted by

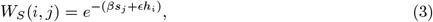

with negative values of *s*_*j*_ and *h*_*i*_ respectively denoting extraction of resources from the host by the symbiont and further benefits provided to the symbiont by the host, and positive values denoting the opposite. Here, *β* and *ϵ* denote selective advantages of incremental negative changes in *s*_*j*_ and *h*_*i*_, respectively, from the symbiont’s perspective. (In the following, we assume positive values for the coefficients *α, β, δ*, and *ϵ*, so that a cooperative symbiont has a positive value of *s*; a change in sign for the coefficients simply shifts the interpretation of the signs on *h* and *s*).

Using 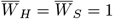 as the benchmark for neutrality (effectively the situation in which the participants cannot evolve to influence each others fitness beyond the obligatory nature of their relationship), this weighting scheme partitions the phenotypic coordinates of the two species into four informative quadrants (Figure 1). For example, again assuming positive selection coefficients (*α, β, δ, ϵ* > 0), positive *h* and *s* implies that *W*_*H*_ > 1 and *W*_*S*_ < 1, indicating that the system has evolved to a point where the host exploits the symbiont to the detriment of the latter. In contrast, negative *h* and *s* implies that *W*_*H*_ < 1 and *W*_*S*_ > 1, indicating that the symbiont exploits the host. Positive *h* and negative *s* raises the possibility of mutual benefits / exploitation, although the degree to which the fitnesses of the participants deviate from the neutral expectation will depend on the relative weightings of the selection coefficients. Likewise, negative *h* and positive *s* raises the possibility of mutual antagonism, again to a degree that depends on the relative strengths of selection.

### Mutational transitions

Individual sites within genotypes of host cells are assumed to mutate with rates *u*_*H*_ for downward (+ → −) mutations and *b*_*H*_*u*_*H*_ for upward mutations over the entire *h* array, with *b*_*H*_ accounting for potential mutational bias. Mutation rates are assumed to be low enough that any genotype can only mutate to adjacent states in a stepwise fashion, although the genotype-wide mutation rates to such states depend on the parental genotypes, which define the number of mutational targets in the + → − vs. + → − directions. For example, letting *i* denote the number of + alleles within a host haplotype, the mutation rates to one step lower and one step higher in the genotypic array are

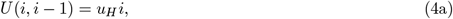

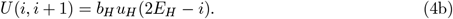

In principle, things are more complicated for the symbiont distribution, as there are multiple symbionts per host cell. However, to reduce the complexity of the model, we assume that host cells are effectively homoplasmic, always containing just a single symbiont type. This is justified by the fact that individual host cells generally contain small to moderate numbers of symbionts, ensuring relatively rapid loss or fixation of newly introduced mutant symbionts (within host cells) on the timescale of evolution of the entire host population. Within a given host cell, establishments of single-step upward or downward transitions from the resident symbiont type then arise with probabilities equal to the product of the rate of introduction of variants by mutation within host cells (assumed to arise in single individuals just once per host generation, owing to mutation rates per individual symbiont ≪ 1) and their probability of internal fixation.

For example, letting *K* be the number of symbionts per host cell, the probability of downward mutations (*s*_*j*_ → *s*_*j*−1_) in the total symbiont population within a host cell is *Kju*_*S*_. The transition rate from type *j* to *j* − 1 is obtained by multiplying *Kju*_*S*_ by the internal fixation probability,

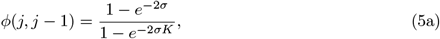

where

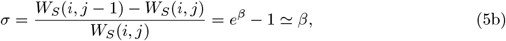

is the within-host cell selective advantage (negative *σ* implies a disadvantage), using Equation 2 to define the fitnesses. The same procedure is used for forward mutations, but with *K*(*E*_*S*_ − *j*)*b*_*S*_*u*_*S*_ for the mutation rate and

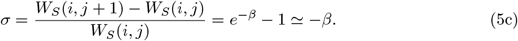

These expressions show that of the parameters in Equation 2, only *β* influences the evolutionary dynamics of the symbiont within a host cell, because all pairs of internally competing symbiont genotypes experience the same host effect. In both cases, if |*Kσ*| ≪ 1, within-cell drift overwhelms selection, the fixation probability ≃ 1*/K*, and the transition probabilities reduce to *ju*_*S*_ or (*E*_*S*_ − *j*)*b*_*S*_*u*_*S*_.

### Symbiont sorting

Generations are discrete, and as the two species are incapable of surviving to reproduce on their own, there are no symbiont-free hosts, and all hosts are assumed to contain a constant number *K* symbionts. However, the possibility of host-symbiont genotype reassortment is allowed for via symbionts released to the environment and restored to a viable state after uptake by an alternative host cell. This is implemented by letting the reassociation rate be *m*, where *m* = 0 means an obligatory association without any possibility of horizontal transfer, and *m* = 1 implies random reassortment each generation such that each host cell acquires a single immigrant symbiont (which might have the same genotype as contained in the recipient host cell) (Figure 2). The coefficient *m* is equal to the probability that a host cell receives an immigrant, which then enters with initial frequency 1*/K* and fixes with a probability that depends on its fitness relative to the resident symbiont, which accounts for the remaining frequency 1 − (1*/K*). The latter fixation probability is again given by Equation 5a, with the selection coefficient depending on the fitness of the migrant relative to the resident symbiont (as in Equations 5b,c but allowing for a wider distribution of differences, as migrants might differ by more than a single mutation). The migrant pool is assumed to be drawn randomly from the entire population of symbionts, with the frequency of each symbiont genotype in the pool being equal to that in the current population. Under this scheme, if the resident symbiont haplotype *j* has frequency *p*_*j*_ over the entire population of host cells, then a fraction *mp*_*j*_ of migrations into host cells containing type *j* symbionts have no consequences, and the remaining fraction *m*(1 − *p*_*j*_) are randomly drawn from the pool of nonself symbiont haplotypes.

**Figure 2.**
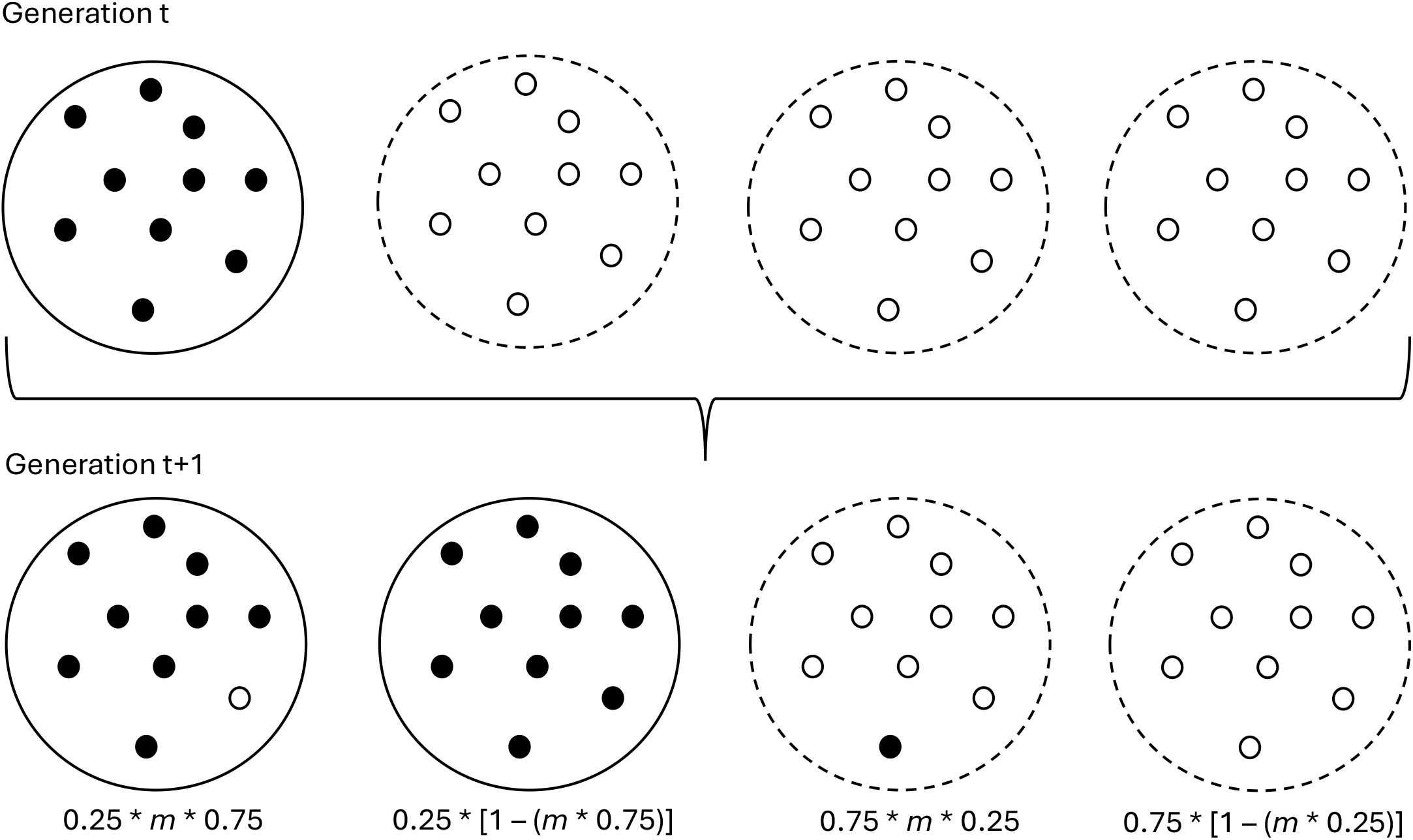
A simple view of the assortment scheme associated with the model. In this example, prior to assortment, there are two cell types, with frequencies 0.25 and 0.75, which also equal the frequencies in the migrant pool of symbionts. The probability of a host cell gaining a single immigrant is *m*, and this creates new configurations if the immigrant genotype differs from the residents. Following such reassortment, intra-host selection operates on the immigrant type (initially in a single copy), leading to either complete loss or fixation, returning all host individuals to homoplasmic states.

### Selection and random genetic drift

Following prior episodes of ressortment and mutation, selection operates on the collection of host-cell/symbiont haplotype combinations by weighting the pre-selection frequencies, *P* ′(*i, j*), by their relative fitnesses,

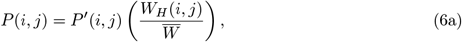

where

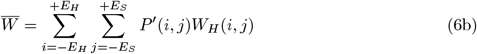

is the mean fitness averaged over the full frequency distribution of host / symbiont genotype combinations following reassortment and mutation but prior to selection.

Random genetic drift is then implemented by multinomial sampling (in a standard Weight-Fisher procedure) based on the post-selection frequencies. The next cycle of reassortment, mutation, selection, and drift then proceeds, with such iterations continuing for long enough time periods to obtain the quasi-steady state genotypic constitution of the population.

## RESULTS

Two levels of selection operate in this system: among symbionts within host cells (via the fixation of new mutations or immigrants resulting from horizontal transfer), and among host cells based on the composite properties of the host and symbiont traits. Thus, the model is set up in a way that allows the system to evolve to a steady state involving conflict and/or cooperation between the two participants. The equilibrium point to which the system evolves is a function of the relative strengths of three types of selection and of the power of random genetic drift at the host and symbiont levels (as noted above, the fourth selection coefficient *ϵ* has no influence on the steady-state solutions, as competing symbionts always vie for fixation within the same host cell). The frequency of horizontal transfer of symbionts among host cells will also be shown to play a central role.

To determine the qualitative outcomes of alternative situations, we systematically ran simulations with *α, β*, and *δ* set to various values, including boundary conditions in which one or two coefficients were set to zero. The parameters for the selection coefficients were typically in the range of 0.0001 to 0.01; symbiont numbers per host cell were *K* = 100, 10^3^ or 10^4^; and host-population sizes ranged from *N* = 10^3^ to 10^7^. This sets up a range of situations in which the strength of selection at the two levels ranges from effective neutrality to very strong relative to the power of drift. Mutation rates per genomic site were typically in the range of 10^−7^ to 10^−5^ per host-cell generation with no directional mutational bias. Finally, purely vertical transmission was modeled with *m* = 0, and ample opportunity for horizontal transmission was provided by letting *m* = 1 (every host cell receives one immigrant symbiont per generation). This being said, for a number of cases outlined below, analytical solutions consistent with the simulation results were obtained in terms of the underlying parameters, reducing the need for assumptions about particular parameter values and rendering the scaling relationships more transparent.

### Cross-species neutrality

In the simplest situation with *α* = 0, the host’s fitness only depends on its own genotypic value, and provided *β* ≠ 0, the symbiont’s fitness depends only on its genotypic state. In this case, although the two species are locked together in an obligate symbiosis by prior historical factors unassociated with the the traits under selection, they are otherwise on independent evolutionary trajectories, both evolving in a direction that favors self but to a degree that depends on the strength of selection relative to that of drift.

For the case in which symbiont transmission is purely vertical (*m* = 0), provided mutation rates are sufficiently low (discussed below), one can anticipate that the genotypic means for both members of the pair can be approximated by the formula of Li (1987) and Bulmer (1991), which predicts the steady-state probability of a + allele under the joint processes of drift, mutation, and selection,

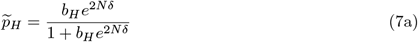

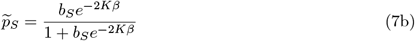

for the host and symbiont respectively, where *b*_*x*_ is the mutation bias in genome *x* (ratio of mutation rates for − to + alleles vs. the reciprocal, with *b*_*x*_ = 1 in the case of no bias). Note that if the absolute value of the product of population size and selection coefficient at a particular level (*Nδ* or *Kβ*) is ≪ 1, drift overwhelms selection and the equilibrium allele frequency will be the neutral expectation *b*_*x*_/(1 + *b*_*x*_). In contrast, if this product is ≫ 1, the favorable allele will be essentially fixed at all times by positive selection. Only when the terms in the exponents have absolute values in the range of 0.1 to 10.0 is there a noticeable effect of population size on the equilibrium allele frequency. In Supplemental Text A, we show that symbiont migration (horizontal transfer) alters Equation 7b to

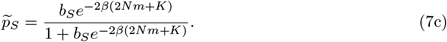

The mean genotypic values are given by

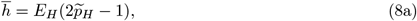

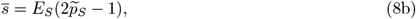

Although Equations (7a-c) are derived under the assumption that the population is not segregating more than one variant at a time (the so-called weak-mutation/strong-selection regime), results from computer simulations are in close accord with the theory under this sequential-fixation regime (Figure 3).

**Figure 3.**
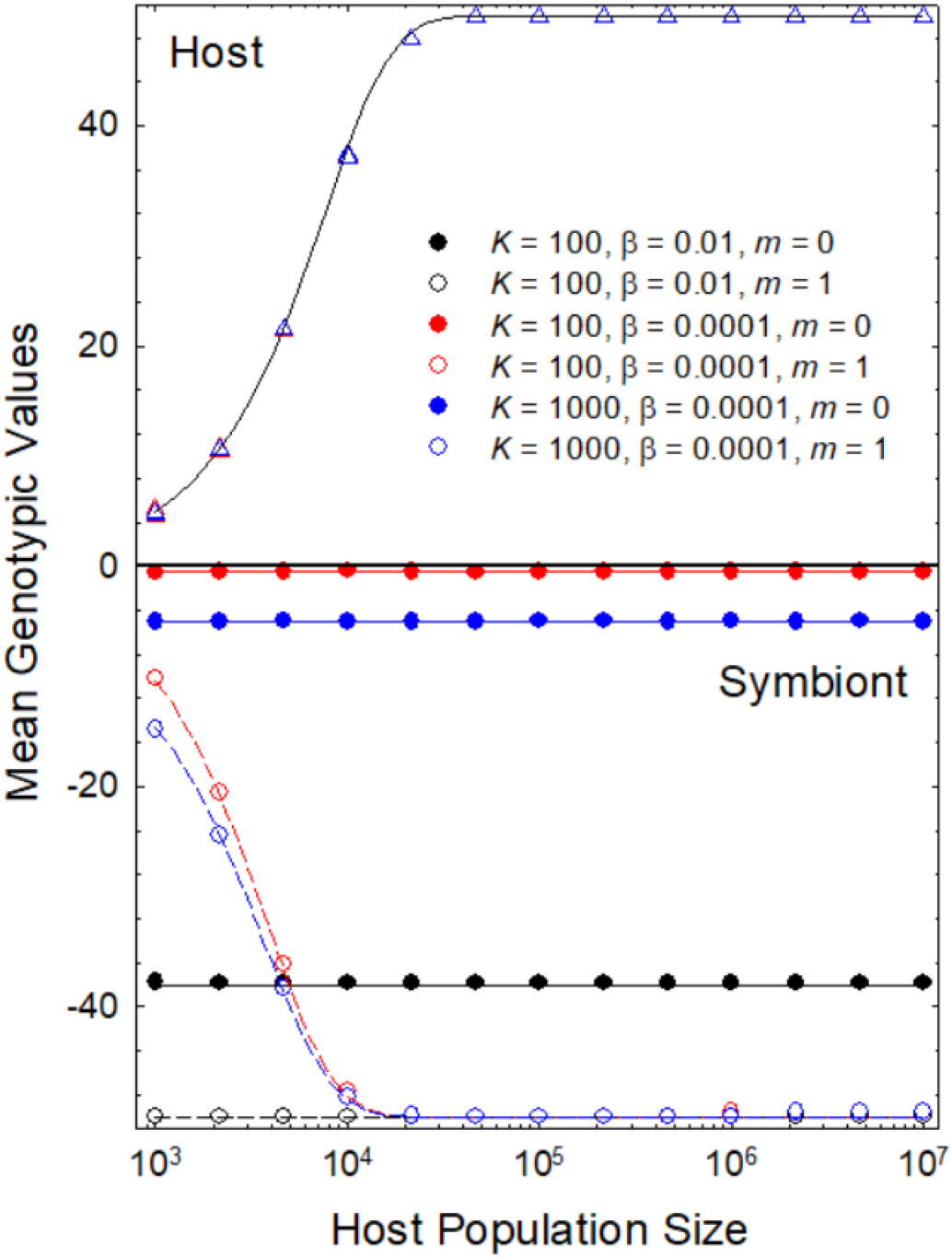
A comparison of the steady-state mean phenotypes of hosts (triangles, upper panel) and symbionts (circles, lower panel) for some cases with and without migration, for the special case in which neither participant influences the fitness of the other (*α* = 0), there is no mutatoin bias (*b*_*H*_ = *b*_*S*_ = 1), and mutation rates at both levels are 10^−7^ per site. The strength of selection on host genotypes is *δ* = 0.0001. The results are in close agreement with the expectations from Equations 7a, 8a (hosts) and 7c, 8b (symbionts), given by the solid and dashed lines. Note that the evolution of the host mean phenotype is completely independent of the symbiont properties, although the reverse is not true.

Several conclusions are immediately apparent from the theory. First, under this sequential-fixation model, the steady-state solutions are completely independent of the mutation rates, although they are influenced by mutation bias. Second, the term *e*^2*Nδ*^ is equal to the ratio of fixation probability of − → + mutations to that in the reciprocal direction at the host level, so *b*_*H*_*e*^2*Nδ*^ is the net pressure of mutation and selection in the direction of + alleles in host genomes. If *e*^2*Nδ*^ ≪ 1, 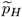 evolves to the neutral expectation *b*_*H*_/(1 + *b*_*H*_), whereas any mutation bias is overwhelmed when *e*^2*Nδ*^ ≫ 1, resulting in 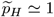 and evolution to the extreme value 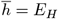.

Similar arguments can be made for the evolution of the symbiont. In the absence of horizontal transfer (*m* = 0), evolution of the symbiont is completely independent of the host population size, and each host cell can be viewed as an isolated island harboring *K* symbionts each generation. However, although the host trait has no influence the fitness of the symbionts, provided there is migration, the host population size does have an indirect effect by modifying the effective population size of the symbiont from *K* to 2*Nm* + *K*. This then passively elevates the probability of fixation of favorable mutations within symbiont genomes by reducing the magnitude of drift at the symbiont level. By this means, a large host population can enable a benign symbiont to evolve to its full potential for extracting benefits from the host. Letting *βK* be the ratio of the strength of within-host selection to drift under pure vertical inheritance, horizontal transfer increases this to *β*(2*Nm* + *K*). Thus, one migrant per generation at the total host-population level (*Nm* = 1) is equivalent to adding 2 to the symbiont effective population size within hosts, so even if *m* ≪ 1, horizontal transfer can greatly influence symbiont evolution if *m > K/N* .

### System entirely symbiont driven

This special case is similar to models involving two levels of selection, explored by many previous investigators (e.g., Luo 2014; Luo and Mattingly 2017), usually with two alleles having conflicting effects at the within-vs. between-group levels, e.g., one allele being selfish at the expense of group benefits and the other being cooperative but disadvantageous at the within-group level. The analogy with the analyses here is that the groups in prior work consist of collections of individuals, whereas here the individual constitutes a group of collective symbionts. Whereas the two alleles under previous models are usually not subject to mutation nor often to drift, our goal is to determine the long-term steady-state distribution of an array of multi-site haplotypes under the joint forces of drift, selection, migration, and reversible mutation.

Letting *δ* = 0, and again assuming *β* > 0, the fitnesses of both symbiont and host are dependent only on the symbiont genotypic value *s*, so the host genotype (from the standpoint of the interaction) is expected to evolve towards the neutral expectation

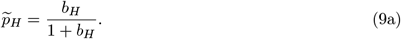

Nonetheless, this situation also sets up a conflict between selection on symbionts at the two levels. At the within-host level, selection favors negative *s* (in conflict with host-cell fitness), which increases symbiont fitness in competitive bouts with each other. In contrast, positive *s* (exploitation of the symbiont) is favored at the host level. Thus, there will be a critical host population size below which the net selective effects are resolved in favor of the symbiont (negative 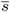) and above which the system favors exploitation by the host (positive 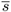).

This behavior results from the different efficiencies of selection operating at the two levels, which can be seen by expanding Equation 7c to include the ratio of strength of positive selection and drift at the host level,

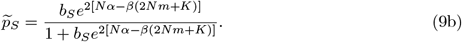

Equation 9b shows that the transition between symbiont- and host-dominated regimes (net negative vs. positive selection on *s*) occurs at the point where the host population size exceeds

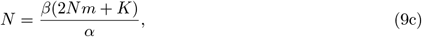

which requires *α* > 2*mβ*. Rearranging to 1 = (*β/α*)[2*m* + (*K/N*)] shows that significant horizontal transfer (*m*), elevated *K/N* (ratio of the within-group vs. total group number), and/or relatively strong selection at the symbiont level (*β/α* > 1) can all tip the balance in favor of exploitation by the symbiont. Near the tipping point, where there is a balance between opposing selection pressures at the host and symbiont levels, 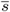 will be free to transiently wander to higher or lower values with fluctuations in host population size. If the efficiency of selection at both levels is high (*αN* and *β*(2*Nm* + *K*) ≫ 1), there can be a nearly stepwise response function, with progression to extreme negative or extreme positive values of 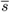 just below and just above the critical point.

Notably, Kimura (1983) considered a model in which an infinite number of groups (in our case, host individuals) harbor finite numbers of group members and derived a tipping point (his Equation 18a) that is identical to Equation 9b, after correcting for haploidy and noting that our *m* is equivalent to his number of migrants per deme. In a somewhat different migration model than employed here, but again focused on conflicts in group selection, Traulsen and Nowak (2006) also arrived at an expression close to Equation 9c, with the key determinants as to whether group members (symbionts, in our case) evolve in the direction of conflict vs. cooperation again being *β/α, K/N*, and *m*.

Some examples are shown in Figure 4, where *βK* = 0.01, 0.1 or 1.0. With high migration (*m* = 1), selection at the level of the well-mixed symbiont population is so efficient that *s* evolves to its extreme negative value to a degree that increases with the host population size. However, in the case of strict vertical inheritance, there is a monotonic increase in *s* to maximally positive values, i.e., from a potentially conflicting situation 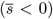 to cooperation at the two levels. With low enough mutation rates to satisfy the conditions of the sequential model, this transition occurs at the point where *αN* = *βK* as expected from Equation 9c.

**Figure 4.**
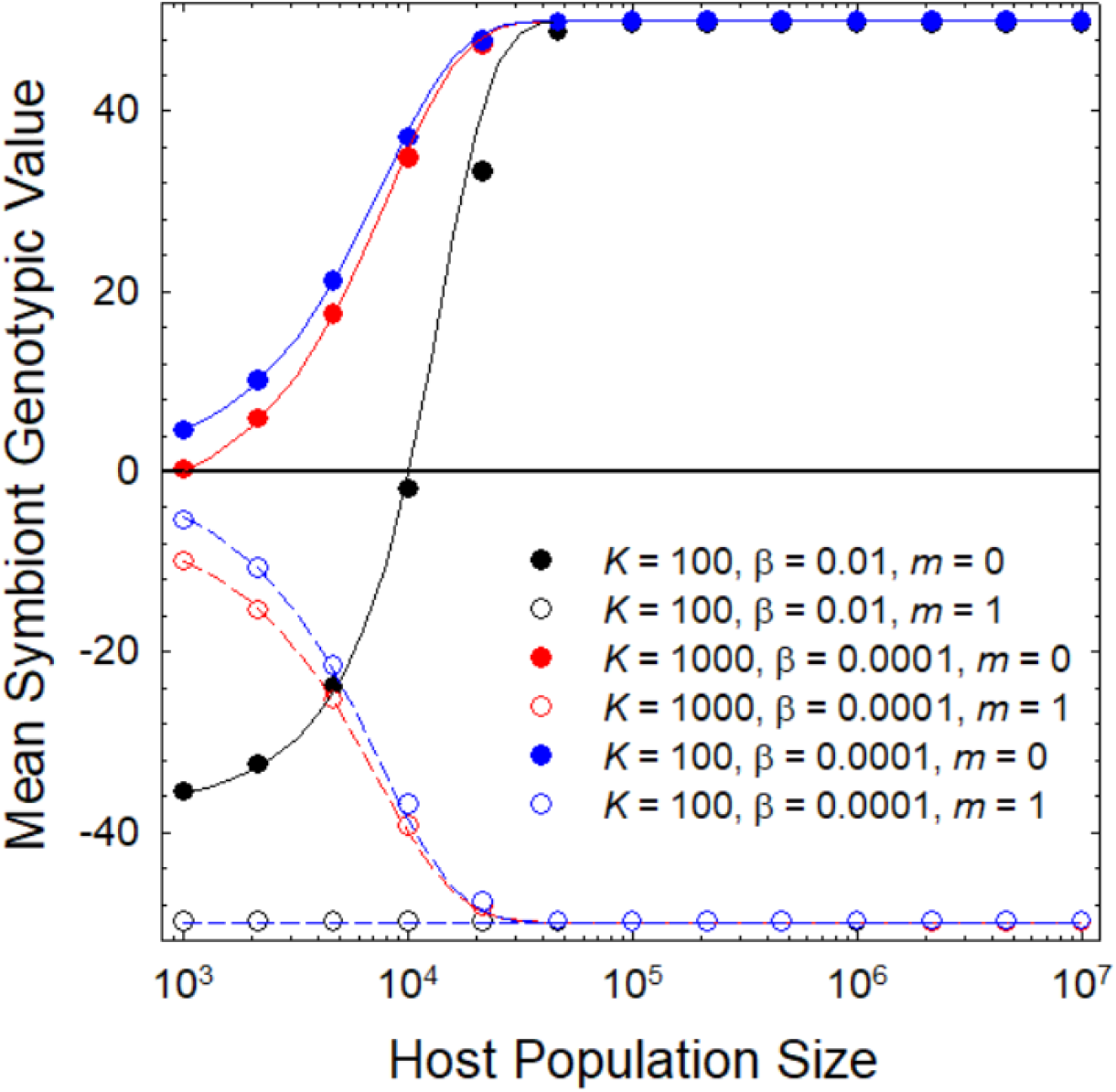
Some examples for the case in which the fitnesses of both participants are only dependent on the genotype of the symbiont; in this case the host genotype evolves to its neutral expectation regardless of host population size (not shown). In all cases for the host, *α* = 0.0001 and *δ* = 0, and both genomes have per-site mutation rates equal to 10^−7^. Data points denote the long-term average values for the symbiont trait, 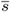, whereas the lines denote the solutions to Equation 9b. Closed points are results for strict vertical inheritance (*m* = 0), whereas open points denote cases in which *m* = 1.

### Joint selection on the host and symbiont traits

With the full selection model (*α, β, δ* > 0), Equation 7a continues to approximate the simulation data for the host population very well, provided mutation rates are low enough to prevent significant selective interference. Not surprisingly, 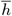 always evolves in a positive direction, asymptotically approaching the maximum value with increasing host population sizes. As noted above, the symbiont has no impact on the evolution of the host trait, as differential fitness among symbionts is independent of the host background, and for the same reason, the symbiont trait continues to behave in accordance with Equation 9b.

The main point again is that strict vertical inheritance facilitates the evolution of a cooperative interaction with increasing host population size, with the transition from a conflicting outcome occurring at the critical *N* denoted above. In contrast, high levels of horizontal transfer encourage the evolution of selfish symbionts. Of particular note here are the cases with horizontal transfer, in which the symbiont trait evolves toward negative values (exploitation of the host), while the host trait evolves towards positive values (exploitation of the symbiont). In many cases, these two opposing changes can be completely balanced, resulting in a stalemate in terms of overall fitness of the host. Consider, for example, Equations 2 and 3 – if the two strengths of selection are equal, but 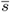 and 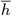 evolve to opposite extremes, the performance of the participant will not have changed, although the structure of the system will have been radically altered relative to the situation before coevolutionary repatterning. That is, the gain that the host acquires by extracting resources from the symbiont is completely balanced by the reciprocal exploitation by the symbiont, and vice versa.

### Symbiont conditioning of the within-host population size

Given that symbionts can influence the fitness of their host cells, it stands to reason that their activity may also have feedback effects on the number of symbionts that can be maintained per host cell. This might occur, for example, if the presence of symbionts influences the size or other physiological features of the host cell, which likely is almost always the case. This, in turn, would influence the effectiveness of selection at the symbiont level by modifying the power of within-host drift. It is unclear whether symbionts that extract resources from the host (*s*_*j*_ < 0) will increase or decrease *K*, for whereas aggressive extraction of host resources might reduce *K, K* might also increase as a consequence of the production of public goods or conditioning the host cell to be larger,.

Likewise, it is unclear whether symbionts that provision the host (*s*_*j*_ > 0) will lead to increased or decreased *K*. The central point is that mutations that generate a boost in *K* can increase the fixation probability of a mutation relative to the expectations under conditions of constant *K*, thereby potentially tipping the balance of evolution at the symbiont level in the direction of host exploitation. On the other hand, if the symbiont population size is kept sufficiently small, its evolutionary potential to exploit the host will be thwarted.

Evaluation of the impact of any such effects requires expressions for the fixation probabilities of mutant alleles in the context of a population size responding to frequency changes within host cells. A general expression has been derived for a wide array of possible functions linking the size of a population with the frequency of a mutant allele (Joshi et al. 2026), including arithmetic, geometric, and harmonic means of allele-specific population sizes, all of which can be applied to the goals herein. However, only in the case of the harmonic-mean model have we been able to obtain closed-form expressions that can be integrated into the formulae developed above (Supplementary Text B), and to enhance transparency we will focus on that here.

Consider the situation in which the expected population sizes of pure symbiont populations scale as

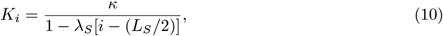

where *i* is the number of + alleles within a symbiont haplotype of length *L*_*S*_, *κ* is the symbiont population size when the haplotype is at the midpoint on the symbiont genotype array (*i* = *L*_*s*_/2), and the weighting factor *λ*_*S*_ must be < 2/*L*_*S*_. Under the harmonic-mean model, during the sojourn of a symbiont mutation or variant immigrant en route to loss or fixation within a host cell, the total number of symbionts within the cell varies in a frequency-dependent manner as

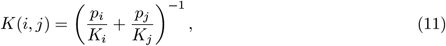

where *p*_*i*_ and *p*_*j*_, which sum to 1.0, are the frequencies of the two types of alleles at any point in time. Equation 9b then expands to

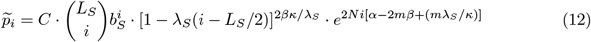

(Supplemental Text B), which converges back to Equation 9b as *λ*_*S*_ → 0.

In the example given (Figure 5), the values of *λ*_*S*_ and *L*_*S*_ yield only a six-fold range of variation in *K*_*i*_ over the full range of symbiont haplotype space, yet there can be up to a two-fold range in the extremes to which the mean symbiont trait can evolve. The effect is necessarily largest at intermediate host population sizes, because at extremely low values of *N*, selection is either completely ineffective or operating at maximum efficiency, respectively. Because the harmonic-mean model is one of the least effective weightings in terms of evolutionary boosts (Joshi et al. 2026), one can anticipate even larger effects with other models (such as the arithmetic or geometric mean), which give more pronounced weightings to rare mutant alleles.

**Figure 5.**
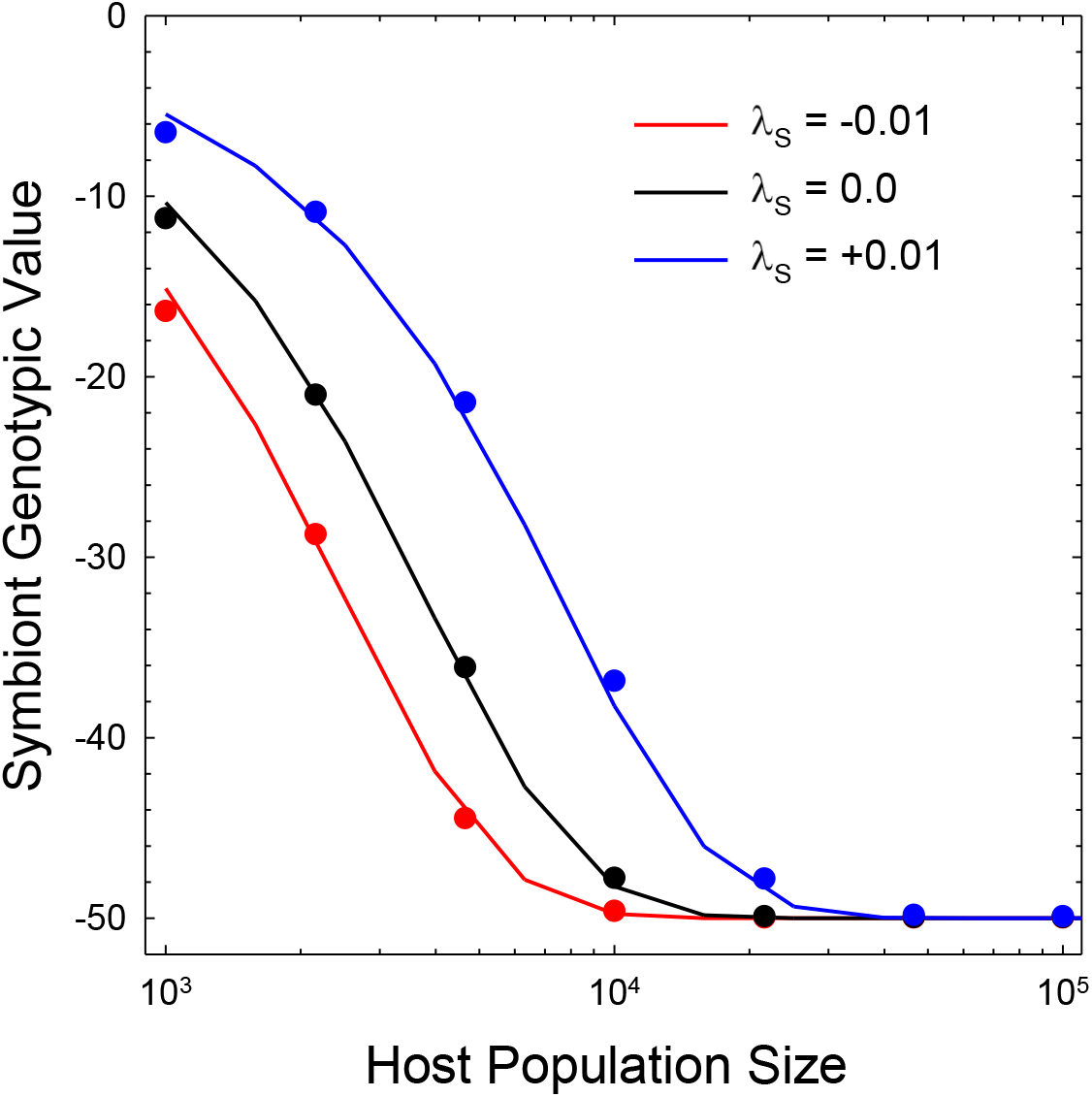
An example of the influence of the influence of symbiont conditioning of the host-cell environment on the evolution of the symbiont trait *s* (more negative values implying a more exploitative symbiont). Here, horizontal transmission occurs at rate *m* = 1, selection on the symbiont trait only occurs through symbiont fitness (*α* = 0), the baseline number of symbionts per host cell is *κ* = 100, symbiont genome length is *L*_*S*_ = 100, and strength of selection on the symbiont trait is *β* = 0.0001 per additional + allele. The lines are obtained by solving Equation 12. The absolute values of *λ*_*S*_ are about half the maximum possible values that can be used for this set of parameters.

### Time-scale difference between hosts and symbionts

It will often be the case that the generation length (cell division time) of the host exceeds that of its symbionts. Indeed, in the preceding development of the model, to minimize the complexity of the derivations and simulations, we assumed that novel symbionts acquired within a host cell by mutation or horizontal transfer are either effectively lost of fixed within the lifespan of the host cell. Adhering to this treatment, but allowing for further extension of the host generation length, we have found that elevating the host generation time by a factor *x* relative to that of the symbionts and reducing the symbiont mutation and migration rates by the same factor *x* (but keeping the host selection coefficients and mutation rates the same over the longer generation length) leaves the steady-state distribution invariant, as this reduction precisely accounts for the additional rounds of mutation and migration events experienced by symbionts within one host generation.

### Selective interference with high symbiont mutation rates

The preceding analytical results were obtained for situations in which new mutations in symbiont and host proceed to loss or fixation independently, with no background interference resulting from other mutations simultaneously in the population. Such interference is not an issue at sufficiently large population sizes, as the participant genomes evolve to their extreme values, with secondary mutations remaining at very low frequencies, nor is it an issue in populations that are sufficiently small that there is a long waiting time between the appearance of consecutive mutations destined to fixation. However, at intermediate population sizes, especially if the number of genomic sites is large and/or the mutation rate per site is high, polymorphic mutations that are simultaneously competing for fixation will interfere with each other’s advancement by selection. This effect can be particularly acute when nucleotide sites are completely linked, as they are here. For example, if for a pair of sites there are two haplotypes simultaneously present in the population, +− and −+, only one can eventually progress to fixation without recombination or secondary mutation.

Although some progress has been made, correcting for interference with analytical expressions has proven difficult, as the process depends on the expected time to fixation of beneficial mutations, the number of sites available for such mutations, and the magnitude of the selective advantages of secondary mutations (Devi et al. 2023; Becher and Charlesworth 2025). The conditions under which the sequential models introduced above are adequate for first-order approximations are presented in Supplemental Text C.

The key point is that selective interference reduces the evolutionary ability of the host to evolve a capacity to exploit its symbiont, i.e., reduces the potential for the evolution of cooperation. Simulation results show that the effects of interference (favoring the evolution of an exploitative symbiont) increase with increasing mutation rates and selection coefficients in the symbiont, as well as with the number of symbionts per host cell (Figure 6). The fact that plausible conditions exist in which selective interference causes a qualitative shift in the nature of the interaction, from cooperation to conflict, supports the contention that differences in the population-genetic environment alone can dictate the long-term outcome of coevolving symbioses.

**Figure 6.**
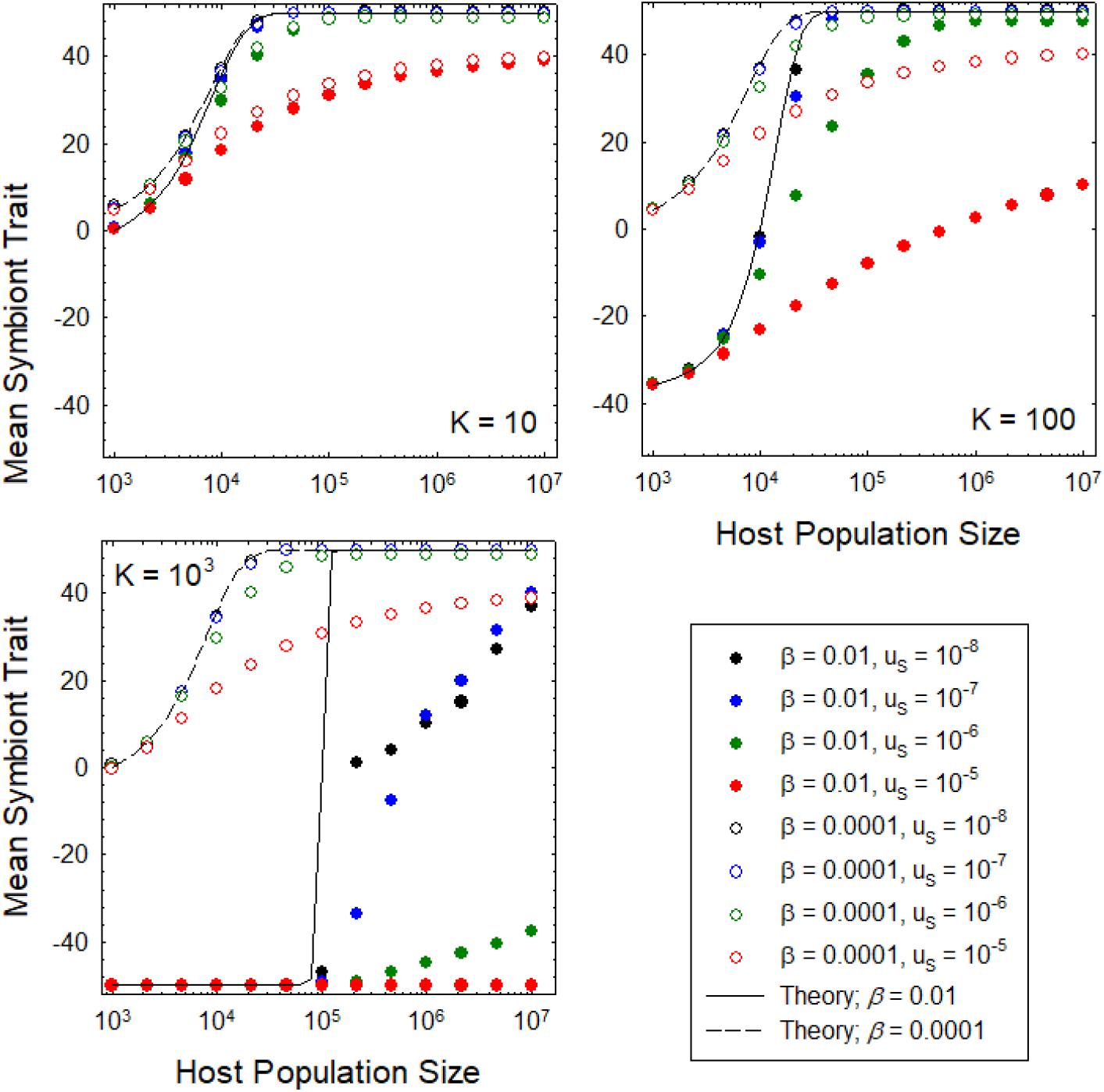
Evolution of the symbiont trait as a function of the number of symbionts per host cell (*K*), the symbiont mutation rate (*u*_*S*_), and the strength of selection on the symbiont for exploitation of the host (*β*). The solid lines give the expectations derived from the sequential model, Equations 8b,9b, so the depression of the results from simulations (closed and open circles) indicates the degree to which selective interference between simultaneously segregating mutations magnifies the ability of the symbiont to evolve an exploitative capacity. Such effects become increasingly more pronounced with increasing *u*_*S*_. More generally, at sufficiently high and sufficiently low host population sizes, the analytical theory coincides perfectly with the simulation results.

## DISCUSSION

Obligate symbioses in eukaryotes always appear at face value to be mutually beneficial, given that the partners have evolved to the point of being inseparable. However, the current day functions of traits need not reflect the conditions at the time of establishment, and demonstrations that one or both codependent members of symbioses have gained anything energetically relative to their condition before the onset of association are lacking. The eukaryotic mitochondrion provides a striking example. Given that a key role of today’s mitochondrion is ATP production, a common view is that the establishment of the mitochondrion yielded an energetic boost to the primordial host cell, and that eukaryotes and all of their downstream embellishments would not have been possible without this profound gift of the hopelessly enslaved victim (Lane and Martin 2010; Lane 2018). However, it now appears that the mitochondrion arose subsequent to the establishment of many of the embellishments of the eukaryotic cell (Pittis and Gabaldón 2016; Hampl et al. 2019; Gabaldón 2021; De Anda et al. 2026; Tobiasson et al. 2026), and the capacity for biomass production is reduced in eukaryotic cells (Lynch and Marinov 2015; Lynch et al. 2023; Lynch 2024).

The model presented here has features in common with many attempts to understand the resolution of conflict vs. cooperation between two participants, including models involving selection within and among groups of individuals of the same species (Kimura 1983; Wilson 1983; Frank 1994; Luo 2014; Cooney et al. 2023). Most such studies start with an assumed biallelic locus with one allele elevating the fitness of individuals within groups but reducing group-level mean fitness and the other having the opposite effects, the goal being to determine the conditions under which selfish vs. altruistic alleles come to dominate the metapopulation.

The difference here is that the underlying evolutionary forces (selection, mutation, drift, and migration) influence traits expressed at two levels, each encoded in different genomes (host and symbiont) locked at the outset into obligatory codependence. The alleles at both levels then evolve over time, eventually reaching a joint steady-state probability distribution that how the fitness of each member of the pair deviates from the null expectation for the case of no cross influence on fitness. The main focus is then on how the population-genetic environment effects the degree to which each member of the pair evolves to contribute in a cooperative vs. exploitative manner.

An early attempt was made by Bergstrom and Lachmann (2003) to think about the evolution of symbioses in this way, but their model was deterministic in nature and avoided explicit genetic features, focusing on a simple binary payoff matrix for the two species. Two studies have taken a more quantitative-genetic approach (Frean and Abraham 2004; O’Brien et al. 2021), but the first of these was confined to the case of regular and complete dissociation of the two participants with host genotypes expressing symbiont genotype preference, whereas the second focused on traits under stabilizing selection and extremely strong mutation.

We find that under a wide variety of conditions, particularly when there is horizontal transfer, the symbiont evolves to be an exploiter of goods produced by the host, and whereas the host also evolves to exploit the symbiont, there is often an evolved balance such that the host fitness is no greater than the null expectation. Thus, our results suggest that many obligatory species interactions that superficially appear to be beneficial mutualisms may instead represent evolutionary stalemates or even net exploitation by the symbiont. The balance is tipped in the favor of exploitation by the symbiont when the within-host group size of symbionts is high, the host population size is small, selection is strong on symbionts relative to hosts, there is horizontal transfer of symbionts, and/or the symbionts have accelerated mutation rates or turnover times relative to the host. We also show that when the symbiont conditions the host cell biology to enhance within-host population sizes, selfish symbionts will profit further from the more efficient selection resulting from the diminished level of within-host drift. Finally, we find that even when conditions favor exploitation of the symbiont by the host (most notably when symbiont inheritance is strictly vertical), selective interference between the effectively linked genomes can substantially increase the host population size necessary to force the evolution of symbiont cooperation. Taken together, these results suggest that the evolutionary enslavement of a symbiont to provide a net benefit to the host requires the confluence of a narrow mix of population-biological features of both participants.

We have pointed out some simple threshold levels for the key parameters that dictate whether the symbiont evolves in the direction of conflict vs. cooperation with the host. These results suggest that only after the efficiency of selection on the host exceeds a certain threshold will the system make a transition from parasitism by the symbiont to one in which the latter provisions the host. If the strength of selection on individual mutations is weak, this transition can be gradual, but with sufficiently strong selection, there can be a nearly switch-like jump from extreme conflict to extreme cooperation. The theory may then help explain the precarious nature of the stability of mutualisms (Sachs and Simms 2006; Sachs et al. 2011; Duron et al. 2015).

While we have explored some of the determinants of outcomes under even the simplest case of obligate symbiosis, aside from extending the theory to facultative symbiosis, several broader questions remain to be resolved. First, although intrinsic conflicts are built into our model in that genome changes in either participant that are beneficial to self are deleterious to the partner, we have assumed a linear relationship between the two effects. One might imagine situations in which the benefits to partners plateau with increasing investment while the costs to self increase linearly (Doebeli and Knowlton 1998), although this will complicate the mathematics, possibly eliminating hope for simple expressions. In addition, the structure of the model that we have developed effectively imposes no influence of the host on competition between symbionts, and therefore no pressure for the host to provision the symbiont – distinct symbiont genotypes vying for fixation within a host cell experience the same host background effect and are only distinguished by their own encoded effects. Thus, although selection at the host level can force the symbionts to evolve towards cooperation, there is no opportunity for the reverse, i.e., for the symbiont to force the evolutionary enslavement of the host. It is difficult to see how the latter effect could be implemented, as selection at the host level ultimately dictates which symbionts survive.

Second, although we have evaluated the consequences of horizontal transfer, we have not considered the symbiont migration rate to be an evolvable trait. Prior work has suggested why the capacity for transfer might be under selection (Frank 1996). One consideration is that migration leads to competition between different symbiont lineages and hence to symbiont phenotypes with more negative effects on the host (as shown here), and that as a consequence, hosts should evolve to restrict horizontal transfer of symbionts to minimize the likelihood of exploitation. The degree to which such second-order (long-term) effects can be promoted over mutations with immediate effects on host / symbiont fitness, and future work should explore the situation in which the degree of transfer is an evolvable feature of traits encoded in both the host and symbiont genomes.

Third, although we have allowed for the possibility that the symbiont can condition host cells in ways that might either increase or reduce the within-host population size (e.g., by altering host-cell size), it is also possible that host population sizes are altered by the presence of symbionts, through direct or indirect ecological effects. For example, with fixed resources available to the host population, a shift to larger host-cell sizes might result in a reduction in host-cell number. This would further reduce the efficiency of selection at the host level, favoring exploitation by the symbiont to an even greater extent. In the Supplemental Text (Equations B11,12), we have sketched out one way in which such effects might be formally incorporated.

Fourth, the analytical theory that we present relies on the sequential model, which assumes that the fixation of consecutive mutations in the two participants proceeds without background interference from different mutations simultaneously segregating in the population at different genomic sites. As pointed out in the results section, violations of this assumption underlying the analytical solutions only reinforce the conclusion that the tendency towards exploitation by symbionts is a likely outcome in most situations. Nonetheless, to improve the quantitative aspects of the overall theory, future work will need to more formally extend to situations in which the assumptions of the sequential model are significantly violated (Supplemental Text C). This too is a difficult area, as even theory for single-species systems requires further development (Devi et al. 2023; Becher and Charlesworth 2025).

Finally, we wish to emphasize that the theory developed above assumes a scenario in which the evolution of symbiosis proceeds in a stable environment, or at least one in which the symbiosis has become an obligately codependent situation prior to the evolutionary resolution of conflict vs. cooperation between the participants. Thus, we are not ruling out the fact that obligate symbioses can sometimes become established by enabling a species pair to inhabit a new ecological niche, as in the case of many sap-feeding insect endosymbioses. However, the downstream evolutionary issues remain, and one can hardly view the expansion of an ancestral cicada to one locked into a 17-year life cycle as an energetic advancement.

## ACKNOWLEDGMENTS

This research was supported by National Institutes of Health award R35-GM122566-01, National Science Foundation award MCB-1518060, and Moore Foundation Grant 735927.

## SUPPLEMENTAL TEXT

### A) Situation in which the symbiont trait only influences symbiont fitness

In this case, the host trait evolves towards its upper limit, to a degree that only depends on the scaled selection coefficient *δN* and the host mutation bias *b*_*H*_ . As the relative fitnesses of alternative symbiont types depend only on the number of + alleles in their genotype (here denoted as *j*), and not on the host-cell background, the symbiont fitness can be written as

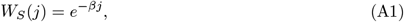

so that the selective difference between any two adjacent trait values is *β*, with reductions in the index *j* being advantageous. We further assume that symbiont mutation rates and population sizes are independent of *j*.

Here, we consider the transition probabilities (*U*) between adjacent symbiont states within individual host cells, which are functions of the mutation rates to destination alleles and their probabilities of fixation. For changes to the next highest index,

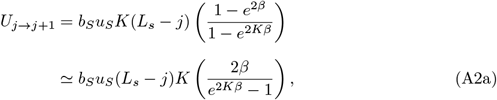

where the final approximation holds for *β* ≪ 1. For changes to the next lowest index,

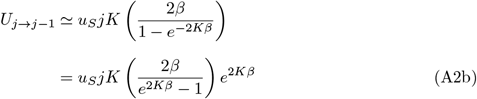

where *L*_*s*_ is the number of genomic sites, *K* is the number of symbionts per host cell, *u*_*S*_ is the mutation rate from + to − alleles, and *b*_*S*_*u*_*S*_ is the reciprocal mutation rate. Thus, these two transition types are equivalent to an effective mutation process with forward rate

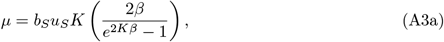

and backward rate *µB* with

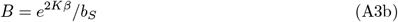

denoting the directional bias. Finally, the rate at which a host cell changes from state *j* to *k* by migration (assumed to be a single immigrant per host cell per migration event) is

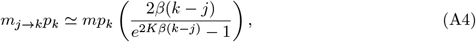

assuming *βL*_*s*_ ≪ 1, where *m* is the frequency of migration events per host cell, and *p*_*k*_ is the frequency of type *k* symbionts across the entire host population (and in the pool of migrants).

Assuming that joint mutation / migration events do not occur, the general recursion equation for the haplotype frequencies of symbionts (assumed to be homoplasmic) is

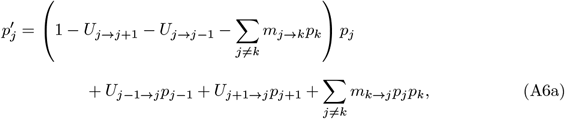

assuming all *U* and *m* terms are ≪ 1, which rearranges to

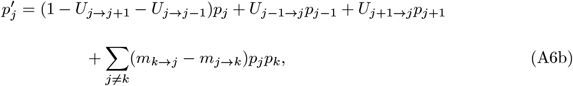

where the two rows, respectively, denote changes due to mutation and migration. Using Equation A4, the migration term simplifies further to

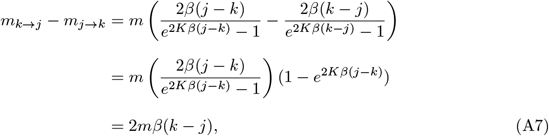

which should be a good approximation provided *βL*_*s*_ ≪ 1. With this transformation, state *j* can be viewed as having an absolute selection coefficient equal to −2*mβj* relative to the benchmark of zero, such that the selective differences between adjacent states are 2*mβ* for any *j* → *j* − 1 and −2*mβ* for any *j* → *j* + 1.

#### Approximation for the small-population-size domain

The preceding analysis reveal a rescaled system of equations with constant effective rates of forward and reverse terms for the combined effects of mutation and within-host selection per site and linear dependence of the fitness differences between haplotypes and their individual states. In the small-population-size regime, the population remains largely monomorphic with rare excursions to adjacent states, initiated by a host cell containing a one-step change in its endosymbionts caused by mutation and within-host selection, with such a variant then having the capacity to spread through the entire host population via the joint process of migratory exchange and within-host selection.

This process has the same form as that for a linear array of nonrecombining haplotypes with *L*_*S*_ biallelic (+/−) sites, with constant rates of mutation per site type and constant strengths of selection between adjacent haplotypes. The balance between the joint forces of mutation, selection, drift, and migration leads to predictions for the steady-state distribution of alternative haplotypes, where *j* indeses the number of + alleles,

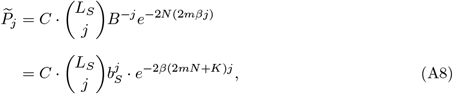

where 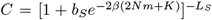 is the normalization constant needed to ensure that the frequencies sum to 1.0 (Foundations 14.2; Lynch 2024).

In this small-population-size domain and with the exponential fitness function, the mean frequency of + alleles (equivalent to the fraction of time spent as +) is unaffected by selective modification at other sites, and is given by the standard Li-Bulmer equation,

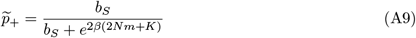

(Li 1987; Bulmer 1991). Equation A9 shows that the total number of migration events per generation (*mN*) boosts the effective population size of the symbiont by 2 (for unclear reasons.)

The mean number of + alleles in the haplotypic array is 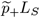, and the mean symbiont state (on the scale of −*E*_*S*_ to +*E*_*S*_) is

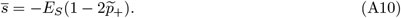

#### Approximation for the large-population-size domain

Here, we assume a host population that is effectively infinite in size, which under the assumption of linkage equilibrium, allows the solution to the deterministic equation

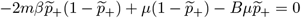

which rearranges to

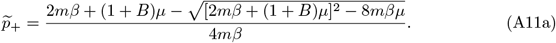

If either *m* ≪ or *m* ≫ *u*_*s*_*K*(1 + *e*^−2*βK*^)/(1 − *e*^−2*βK*^) or *m* ≪ *u*_*s*_*K*(1 + *e*^−2*βK*^)/(1 − *e*^−2*βK*^), then

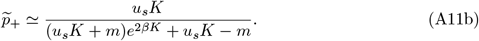

### B) Situation in which the symbiont trait also influences symbiont population size

Here, we attempt to generalize the preceding results to allow for the possibility that symbiont genotypes also influence the within-host population size in a frequency-dependent manner. Letting the population sizes of pure symbiont populations be

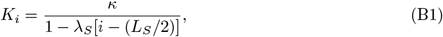

where *i* is the number of + alleles within the haplotype of length 2*E*_*s*_, the weighting factor |*λ*_*S*_| must be < 1*/E*_*S*_, and *κ* is the benchmark symbiont population size when the haplotype is at the midpoint on the symbiont genotype array (*i* = *L*_*s*_/2). For |*λ*_*S*_| ≪ 1*/E*_*S*_, 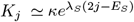. Although other functions for *K*_*i*_ can be imagined, this particular function proves useful mathematically in the derivations below.

A fairly general model for the response of the total intrahost population size for a mixed pair of symbionts of types *j* and *k*, with respective frequencies *p*_*j*_ and *p*_*k*_ (which sum to 1.0, under the assumption of no more than two symbiont types per host cell at any point in time) is given by

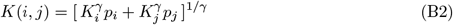

(Joshi et al. 2027). If *γ* = 1, *K* is simply the weighted arithmetic mean of the two haplotype-specific population sizes, whereas *γ* = −1 implies a harmonic mean, *γ* = 2 is a root mean-squared model, and *γ* → 0 yields a geometric mean. This function is required to determine the fixation probabilities of newly arisen mutations of type *j* invading a type *i* population, and changing its frequency in doing so, and vice versa.

An approximate solution to the steady-state distribution of symbiont haplotypes under the small-population-size regime, where transitions between states occur by single-step mutations only, can be obtained by noting that under detailed balance for all adjacent pairs of haplotypes,

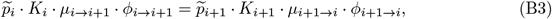

where *µ*_*i*→*i*+1_ and *µ*_*i*+1→*i*_ denote forward and reverse mutation rates, and *φ*_*i*→*i*+1_ and *φ*_*i*+1→*i*_ are the fixation probabilities for forward and reverse mutations. The solution to the full set of equations requires expressions for the ratios of fixation probabilities *φ*_*i*→*i*+1_*/φ*_*i*+1→*i*_, which have the simple form of *e*^2*Kβ*^ (see above) when *K* is constant, but are otherwise more complicated. Although a general expression for the fixation-probability formula has been obtained for the case of frequency-dependent change in *K* (Equation 3a in Joshi et al. 2027), even the expressions for special cases are complicated and nontransparent, as they contain terms in the form of gamma functions, error functions, and exponential integral functions.

#### Harmonic-mean model

Some progress is possible for the case in which *γ* = −1,

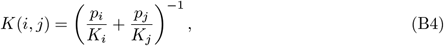

where *p*_*i*_ + *p*_*j*_ = 1, and in the excursion of any mutant allele, the allele frequencies are stochastic variables en route to allele loss or fixation, and

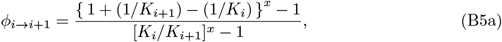

is the probability of fixation of haplotype *i* + 1 on an *i* background (for which there is a selective disadvantage of *i* + 1 relative to *i* of *β*), with

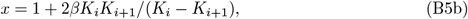

and *φ*_*i*+1→*i*_ obtained by reversing the subscripts in Equation B4, and changing the sign of *β*.

Assuming *K*_*i*_, *K*_*i*+1_ ≫ 1 and using (1 + 1*/K*)^*a*^ ≃ 1 + (*a/K*), leads to

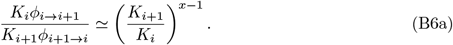

Using Equation B1 yields the recursion equation,

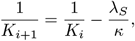

reducing things further to

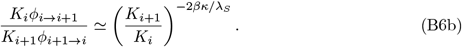

Returning to Equation B3, and noting that *µ*_*i*→*i*+1_*/µ*_*i*+1→*i*_ = (*L*_*S*_ − *i*)*b*_*S*_/(*i* + 1), where *L*_*S*_ is the haplotype length, we obtain

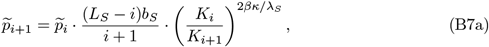

which after applying Equation B1 implies

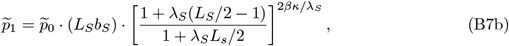

and recursion leads to the general formula

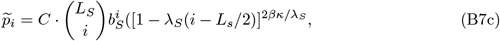

where *C* is the normalization constant that ensures that the frequencies sum to 1.0. In the limit, as *λ*_*S*_ → 0 (a constant population size *K*), the term to the right converges on exp[−2*βK*(*i* − *L*_*S*_/2)], and absorbing the constant part containing *L*_*S*_/2 into the normalization constant yields

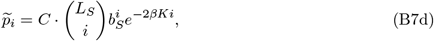

the standard expression for a linear array of alleles separated by constant strength of selection in a population of constant size (Equation A8). Equations B7c,d neatly separate the steady-state distribution into components due to mutation alone (the binomial neutral expectation) and its modification by selection.

Equation B7b assumes strict vertical inheritance of symbionts, but can be generalized to include migration in the following way. Using the same approach as in Equation A7, but with fixation probabilities appropriate for the harmonic-mean value, and assuming *K*_*i*_, *K*_*j*_ ≫ 1, after some algebra,

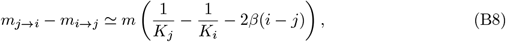

leading to the conclusion that migration leads haplotype *i* to have a net selective effect of

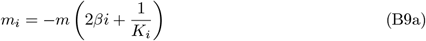

with application of Equation B1 leading to

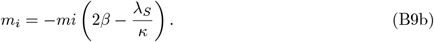

Equation B7c can then be expanded to include the selective effects associated with migration

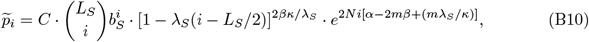

where we have included the term to account for selection at the host-cell level (here assuming that host population size *N* remains constant). Except for the inclusion of selection at the host level, reduces to Equation A8 with constant symbiont population size.

Finally, if the host population size is influenced by the symbiont haplotype, using a function of similar form for the symbiont population size,

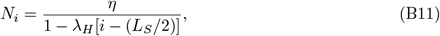

similar steps lead to

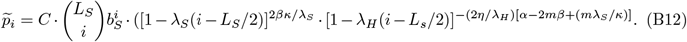

### C) Limits on the domain for the sequential model

The sequential-fixation regime can be approximated by requiring the time to fixation to be less than the time to arrival of next mutation destined to fix. Because the time to arrival of a deleterious mutation destined to fix is generally much greater than that of a beneficial mutation (except in the neutral regime), we focus on the time to arrival of beneficial mutations destined to fix. Under sufficiently strong selection (*Ns*_*b*_ ≫ 1), where *s*_*b*_ is the selective advantage of the beneficial relative to the deleterious allele, the time to fixation ≃ 2 ln(*N*)*/s*_*b*_, and the probability of fixation ≃ 2*s*_*b*_. Thus, the condition for sequential fixation is,

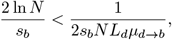

where *L*_*d*_ is the average number of sites with deleterious mutations in steady state, and *µ*_*d*→*b*_ is the mutation rate per site from the deleterious to beneficial state. Rearranging,

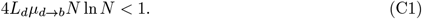

#### Application to two levels of selection

In the system studied here, for the host population, *s*_*b*_ = *δ*, and *L*_*d*_ can be found from the Li-Bulmer solution,

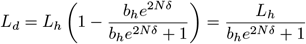

Thus, the condition is,

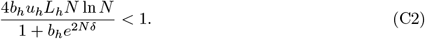

For the symbiont population, following from the text, the net selection coefficient, *s*′ = *α* − 2*mβ*, can be positive or negative, and the identity of the beneficial mutation changes accordingly. The mutation rate towards beneficial mutation also changes,

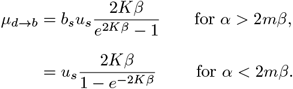

The expected number of sites with deleterious mutations is now,

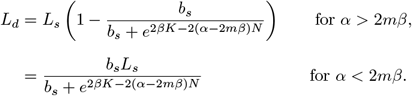

Thus, the conditions necessary to satisfy the sequential model become,

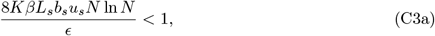

where,

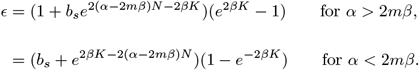

Because we require an overall situation where sequential fixation applies to both host and symbiont traits, we require that Equations 2 and 3 hold simultaneously.

